# Metatranscriptome profiling of the dynamic transcription of mRNA and sRNA of a probiotic *Lactobacillus* strain in human gut

**DOI:** 10.1101/442673

**Authors:** Jicheng Wang, Zhihong Sun, Jianmin Qiao, Dong Chen, Chao Cheng, Xiaotian Luo, Jia Ding, Jiachao Zhang, Qiangchuan Hou, Yi Zhang, Heping Zhang

## Abstract

Metatranscriptomic sequencing has recently been applied to study how pathogens and probiotics affect human gastrointestinal (GI) tract microbiota, which provides new insights into their mechanisms of action. In this study, metatranscriptomic sequencing was applied to deduce the *in vivo* expression patterns of an ingested *Lactobacillus casei* strain, which was compared with its *in vitro* growth transcriptomes. Extraction of the strain-specific reads revealed that transcripts from the ingested *L. casei* were increased, while those from the resident *L. paracasei* strains remained unchanged. Mapping of all metatranscriptomic reads and transcriptomic reads to *L. casei* genome showed that gene expression *in vitro* and *in vivo* differed dramatically. About 39% (1163) mRNAs and 45% (93) sRNAs of *L. casei* well-expressed were repressed after ingested into human gut. Expression of ABC transporter genes and amino acid metabolism genes was induced at day-14 of ingestion; and genes for sugar and SCFA metabolisms were activated at day-28 of ingestion. Moreover, expression of sRNAs specific to the *in vitro* log phase was more likely to be activated in human gut. Expression of rli28c sRNA with peaked expression during the *in vitro* stationary phase was also activated in human gut; this sRNA repressed *L. casei* growth and lactic acid production *in vitro*. These findings implicate that the ingested *L. casei* might have to successfully change its transcription patterns to survive in human gut, and the time-dependent activation patterns indicate a highly dynamic cross-talk between the probiotic and human gut including its microbe community.

**Importance:** Probiotic bacteria are important in food industry and as model microorganisms in understanding bacterial gene regulation. Although probiotic functions and mechanisms in human gastrointestinal tract are linked to the unique probiotic gene expression, it remains elusive how transcription of probiotic bacteria is dynamically regulated after being ingested. Previous study of probiotic gene expression in human fecal samples has been restricted due to its low abundance and the presence of of closely related species. In this study, we took the advantage of the good depth of metatranscriptomic sequencing reads and developed a strain-specific read analysis method to discriminate the transcription of the probiotic *Lactobacillus casei* and those of its resident relatives. This approach and additional bioinformatics analysis allowed the first study of the dynamic transcriptome profiles of probiotic *L casei in vivo*. The novel findings indicate a highly regulated repression and dynamic activation of probiotic genome in human GI tract.

## INTRODUCTION

Microbial communities form an intimate and beneficial association with human gastrointestinal (GI) tract (1-3). Human gut microbiota is of great significance in defending human diseases (4-7). Except for occasional invasion by pathogens, the unique gut microbial ecosystem is continuously exposed to transient microbes originated from diet; while diet can rapidly alter the gut microbial ecosystem (3, 8-10). Metagenomic and metatranscriptomic approaches have been recently emerged as a powerful way to study the impact of pathogens and diet on modulating the composition of human gut microbiota (11-13). However, it remains unclear how the transcriptomes of pathogens and transiet microbes change after entering into the human GI tract. Probiotic microorganisms are generally part of our transient microbiome, which commonly include bacterial strains in the genus *Lactobacillus*, *Bifidobacterium*, *Enterococcus*, and yeasts such as *S. boulardii* (14, 15). After it was coined in 1965, probiotic bacteria have been extensively studied for its wide utilization in dairy foods (16) and prophylaxis and control of a number of disease (17-19), which are primarily focused on their fate, activity and impact on the human gut microbiota (14, 20, 21). Probiotics have been reported to benefit human health in different ways. Probiotics’ capability of rapidly metabolizing some carbohydrates to lactic acid, acetic acid or propionic acid may influence dietary carbohydrate degradation and alter metabolic output, for example, production of short chain fatty acids (SCFA) such as butyrate (14, 22, 23). Many probiotic can establish colonization resistance and competitive exclusion of pathogens (24). Some probiotics are reported to stimulate the human immune response (25-28). However, molecular mechanisms explaining these functions remain largely elusive. Interestingly, a metatranscriptomic study revealed an elevated expression of genes encoding enzymes for carbohydrate utilization in the mouse gut microbiota (29). It should be important to further study who express these probiotic function-related genes, the probiotic bacteria or certain resident microbes?

In general, the distinct probiotic functions and mechanisms should be linked to the gene expression from probiotic microorganisms. Study of the probiotic gene expression in the complicate gut microbe community using traditional methods has been prohibited both by its low abundance and by the presence of closely related species. A recent study has mapped metatranscriptomic reads obtained from elder volunteer fecal samples onto the the probiotic *L. rhamnosus GG ATCC 53103*, showing a good expression of LGG at 28-day of ingestion in a some elders (30). This report promoted us to explore the possibility of using metatranscriptomic reads to study the dynamic of probiotic transcription in human gut.

In this study, we took the advantage of the good depth of metatranscriptomic sequencing reads obtained from fecal samples of healthy young volunteers before and during probiotic ingestion, and extracted strain-specific reads to discriminate the transcription of the probiotic *L. casei* and those of the resident *L. casei*/*paracasei* strains. Strain-specific read analysis showed that transcription of the probiotic *L. casei* was increased while those of its resident relatives remained unchanged. We further showed that transcriptome profiles of the resident *L. casei*/*paracasei* strains and ingested *L. casei* Zhang in human gut were strikingly different. The difference between all *in vivo* transcriptome profiles and those of *in vitro* samples was much more pronounced, and expression of about 40% of mRNAs and sRNA was repressed after being ingested. We observed activation of ABC reporters might be required for probiotic survival during the early stage of ingestion, and genes for sugar and SCFA metabolisms were activated during the later stage of probiotic ingestion. These novel findings underline a highly regulated repression and activation of probiotic genome after being ingested into human GI tract.

## Results

### Experimental design for studying the *in vivo* transcription of an ingested probiotic bacteria

In this study, we used *L. casei* Zhang as a model to study the *in vivo* transcription dynamics of ingested probiotics (Figure 1a). We collected metatranscriptomic reads from the fecal samples taken from six healthy young volunteers (20 to 30 years old, three male and three female) in an open-label clinical trial. The fecal samples were taken on day 0 prior to the consumption and on day 14 and 28 after consumption. Metatranscriptomic cDNA libraries were constructed by respective extraction of RNA from the 18 samples. As controls, we obtained three replicated transcriptomes of *L. casei* Zhang in tablet form prior to the ingestion (Table S1). In order to assess the growth condition of the probiotic in gut microbial community, we additionally sequenced the transcriptomes of *L. casei* cells growing *in vitro* at the lag, log, stationary and death phases (Table S1).

**Figure 1.**
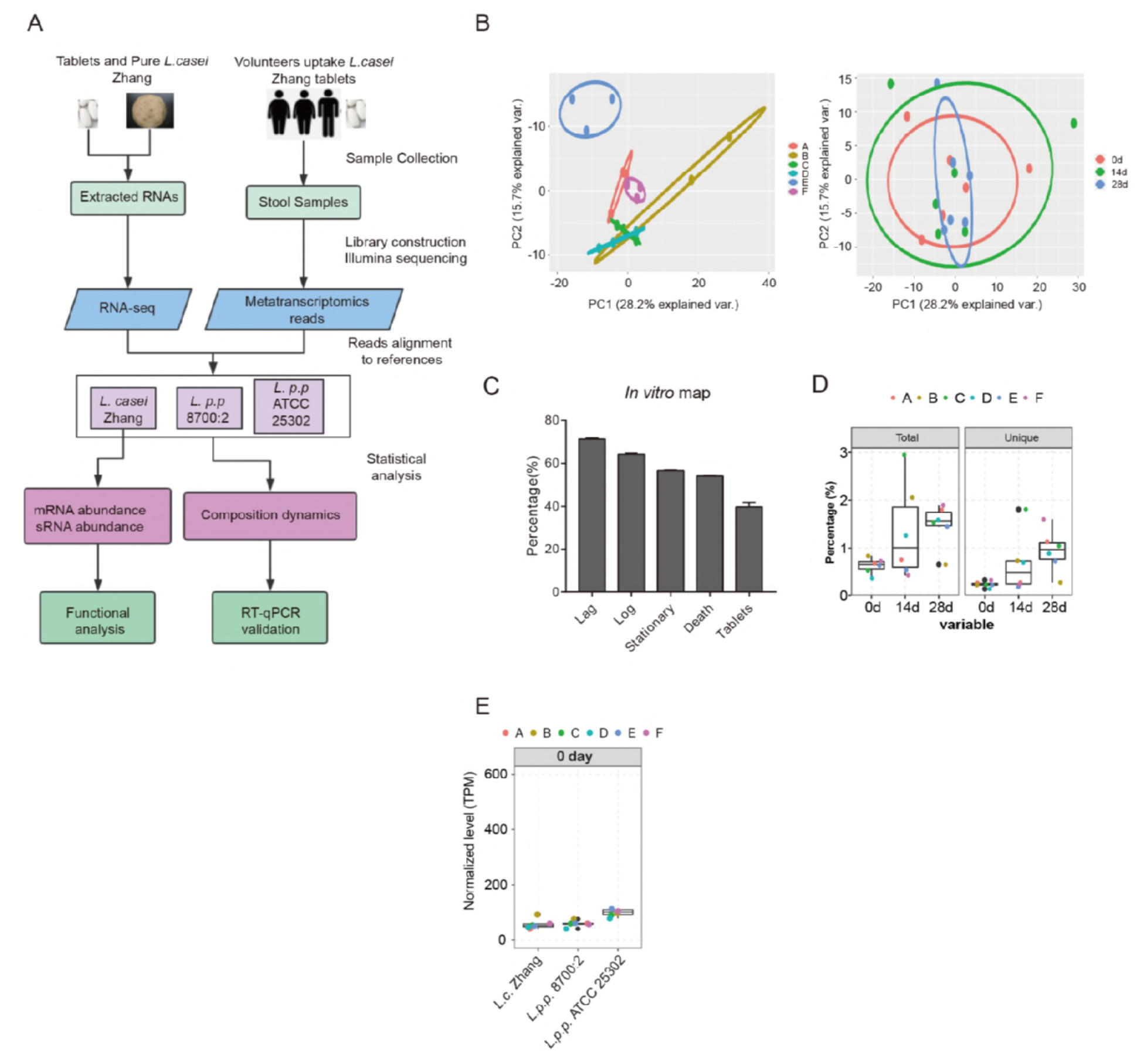
Experimental design and the transcriptional profile of an ingested probiotic bacteria. (A) The flow diagram of this study. Firstly, we took stool samples from six volunteers ingesting *L. casei* Zhang tablets and sequenced the metatranscriptomic reads. In vitro samples were also used to construct RNA-seq libraries, including tablets and cultured pure *L. casei* Zhang. Then we aligned the filtered reads to the reference genomes as well as databases, and calculated the composition dynamics of corresponding species, as well as gene expression abundance. Last, we validated the results by qPCR methods using the original stool samples. (B) PCA analysis showing a large inter-individual variation among all 18 metatranscriptomic samples. The samples were separated by individual classification (left) or by temporal classification (right). (C) Bar plot showing the mapping percentage of *in vitro* samples by aligning the RNA-seq reads to the *L. casei* Zhang genome sequence. (D) Box plot showing the mapping percentage of *in vivo* samples by aligning the RNA-seq reads to the *L. casei* Zhang genome sequence. Left panel was the total mapped reads, and right was the uniquely mapped reads. Dots in each box represent six volunteers. (E) Box plot showing the mapping percentage of *in vivo* samples by aligning the RNA-seq reads to three closely related strains. Samples at 0 day were chosen for representation.

Metatranscriptomic studies of the transcriptional response of gut microbiota in healthy human to the *Lactobacillus* probiotic consumption resulted in controversial observations. One study shows that variation among persons was the biggest reason of transcriptome variation (31), while another study suggests that the transcriptional response of gut microbiota was modulated by probiotic treatment (32). We explored how *L. casei* Zhang affected the transcription/function of our volunteers’ gut microbiotas by analyzing the metatranscriptomic data obtained from the same fecal samples as those of the metagenomic data. Expression correlation analysis showed a large inter-individual variation among metatranscriptomes (Figure 1b). The probiotic-induced change of metatranscriptomes was much smaller than the inter-individual variations (Figure 1b), confirming the lack of a global transcriptional response by probiotic ingestion (31).

Using the genome of *L. casei* Zhang as the reference sequence, we mapped the transcriptome reads from all *in vitro* cultured *L. casei* Zhang, as well as the *in vivo* metatranscriptome and metagenomic samples. About 37.61%-71.62% of transcriptomic reads from *in vitro* cultured *L. casei* Zhang were mapped. The mapping efficiency varied with the culture condition, with the log-phase samples showing the highest efficiency and tablet samples showing the lowest efficiency (Figure 1c). As high as a few percent of *in vivo* metatranscriptomic reads were mapped onto the genome sequence of *L. casei* Zhang after ingestion (Figure 1d). It is shown that mapping results of metatranscriptomic reads were increased in an ingestion time-dependent manner (Figure 1d). Further analysis showed that the base level of mapped reads could be resulted from the presence of closely related strains of *L. casei* Zhang, particularly *L. paracasei* subsp. *paracasei* 8700:2, *L. paracasei* subsp. *paracasei* ATCC 25302 (Figure 1e).

### Transcripts from the ingested *L. casei* Zhang increase significantly while those from the resident *L. casei/paracasei* strains remain unchanged

To further distinguish the transcriptional response of resident *L. casei*/*paracasei* to *L. casei* Zhang ingestion, we only extracted the reads mapped to *L. casei* Zhang, *L. paracasei* subsp. *paracasei* 8700:2 and *L. paracasei* subsp. *paracasei* ATCC 25302, which resulted in strain-specific reads. Plot of the strain-specific reads showed that *L. paracasei subsp. paracasei* ATCC 25302 strain was the most-enriched strain prior to *L. casei* Zhang ingestion. It was interesting to find that, although not dominant, a significant fraction of reads specifically mapped to *L. casei* Zhang in each individual, indicating that *L. casei* Zhang is one *Lactobacillus* strain well adapted to human gut microbe community (Figure 2a).

The overall transcripts from *L. casei* Zhang were increased with the ingestion time, which was anticipated from the successful ingestion of exogenous *L. casei* Zhang. In contrast, the overall transcripts from the other two resident *L. casei*/*paracasei* did not change during the course of investigation (Figure 2b). Consequently, as the fraction of reads mapped to *L. casei* Zhang increased significantly during the course of probiotic ingestion, the resident *L. casei/paracasei* strains decreased (Figure 2c). We therefore concluded that the metatranscriptomic reads mapped to *L. casei* Zhang were dominantly expressed from the ingested *L. casei* at day-14 and day-28.

**Figure 2.**
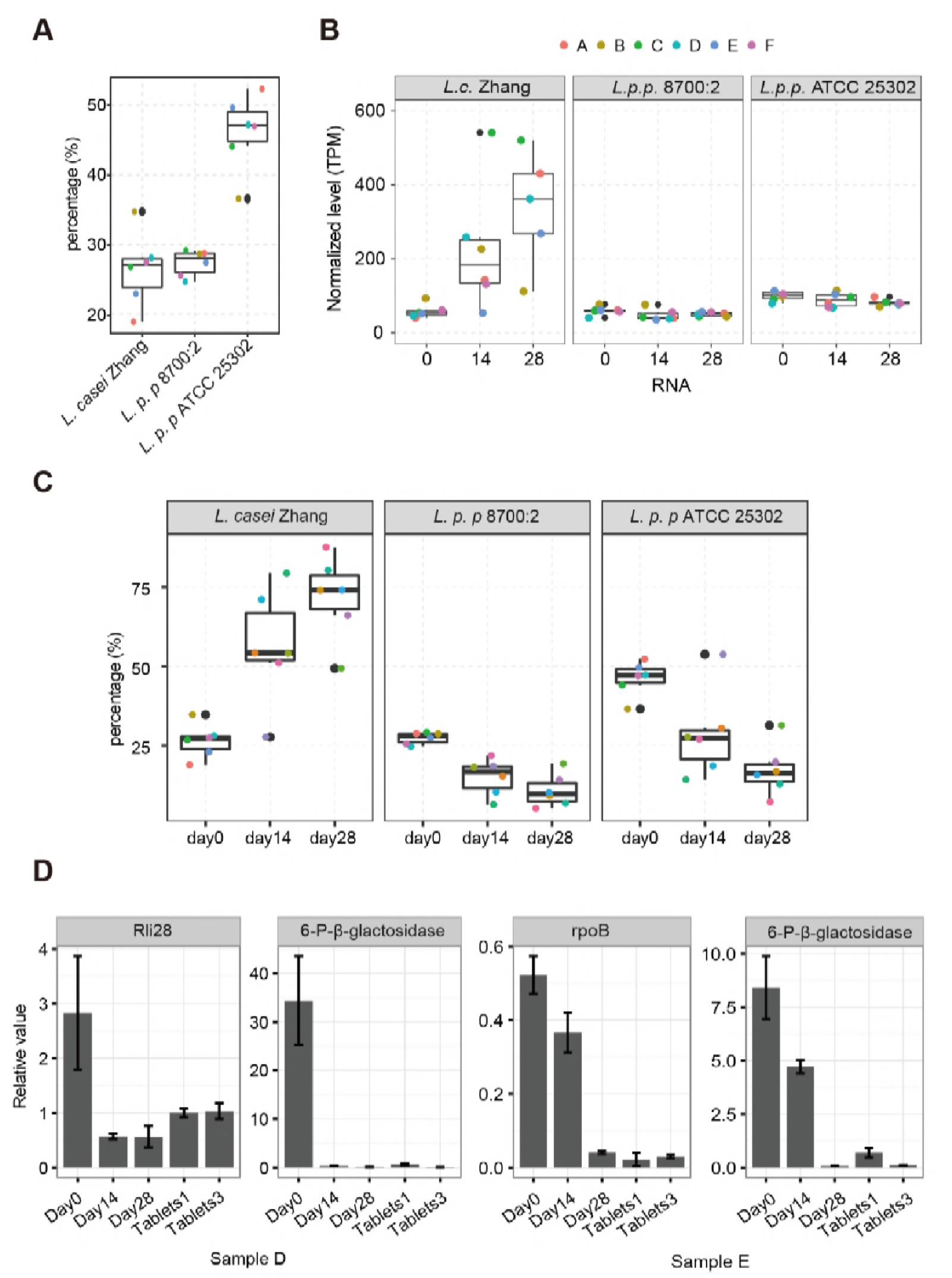
Transcripts from the ingested *L. casei* Zhang increase significantly after ingestion. **(A)** Box plot showing the percentage of mapped reads in each species to the total mapped reads in these three species. Dots in each box represent six volunteers. **(B)** Box plot showing the mapping percentage of *in vivo* samples by aligning the RNA-seq reads to three closely related strains. Samples at three time points were chosen for representation. **(C)** Box plot showing the mapping percentage of reads that were aligned to the combined genome sequences of these three species. **(D)** Bar plot for the RNA/DNA ratio of all these genes by PCR showing the high level expression in the resident *L. casei* Zhang than the ingested grown in in vivo or in vitro (tablets s1 and ts3 samples).

We then selected several genes for absolute quantitative PCR analysis, including β-galactosidase for galactose metabolism and RNA polymerase β subunit (rpoB). Primer for these genes were designed to detect both the ingested and resident *L. casei*/*paracasei*. Figure 2d shows that all of these genes were higher for the resident *L. casei*/*paracasei* (day-0) and lower after the ingested. Their levels in tablets were low and similar as those from human gut after being ingested. Therefore, the transcription pattern of the ingested *L. casei* inherits some of its *in vitro* growth patterns.

Taken together, our mapping results reflected a combined transcription from both the ingested and resident *L. casei/paracasei* strains. We decided to use all reads mapped onto the genome of *L casei* Zhang for the following analysis given the following two reasons. First, the specific reads represented only a very small fraction of the transcriptome and genes. Therefore, gene expression level is hardly to be calculated from the stain-specific reads. Second, transcription from the resident *L. casei/paracasei* strains was repressed after probiotic *L casei* ingestion. Therefore, the observed increase was primarily from *L casei* Zhang.

### Gene expression patterns of *L. casei/paracasei in vivo* are distinct from those *in vitro*

To explore the *in vivo* transcription profiles of different states of the ingested *L. casei*/*paracasei*, we first compared the expression of *L. casei* Zhang among all the *in vivo* and *in vitro* samples. Principal Component Analysis (PCA) of the expression profiles revealed that all *in vitro* transcriptomes were well separated from the *in vivo* transcriptomes by the first component, which indicated a substantial difference between the *in vivo* and *in vitro* transcriptions of *L. casei*/*paracasei* (Figure 3). For the *in vivo* samples, day-28 and day-0 samples were well separated by their transcription profiles (first and second components). However, the patterns of *L. casei*/*paracasei* transcription in the day-14 samples were highly divergent, probably indicating a highly dynamic stage for *L. casei*/*paracasei* transcriptomes responding to the newly ingested *L. casei* Zhang.

**Figure 3.**
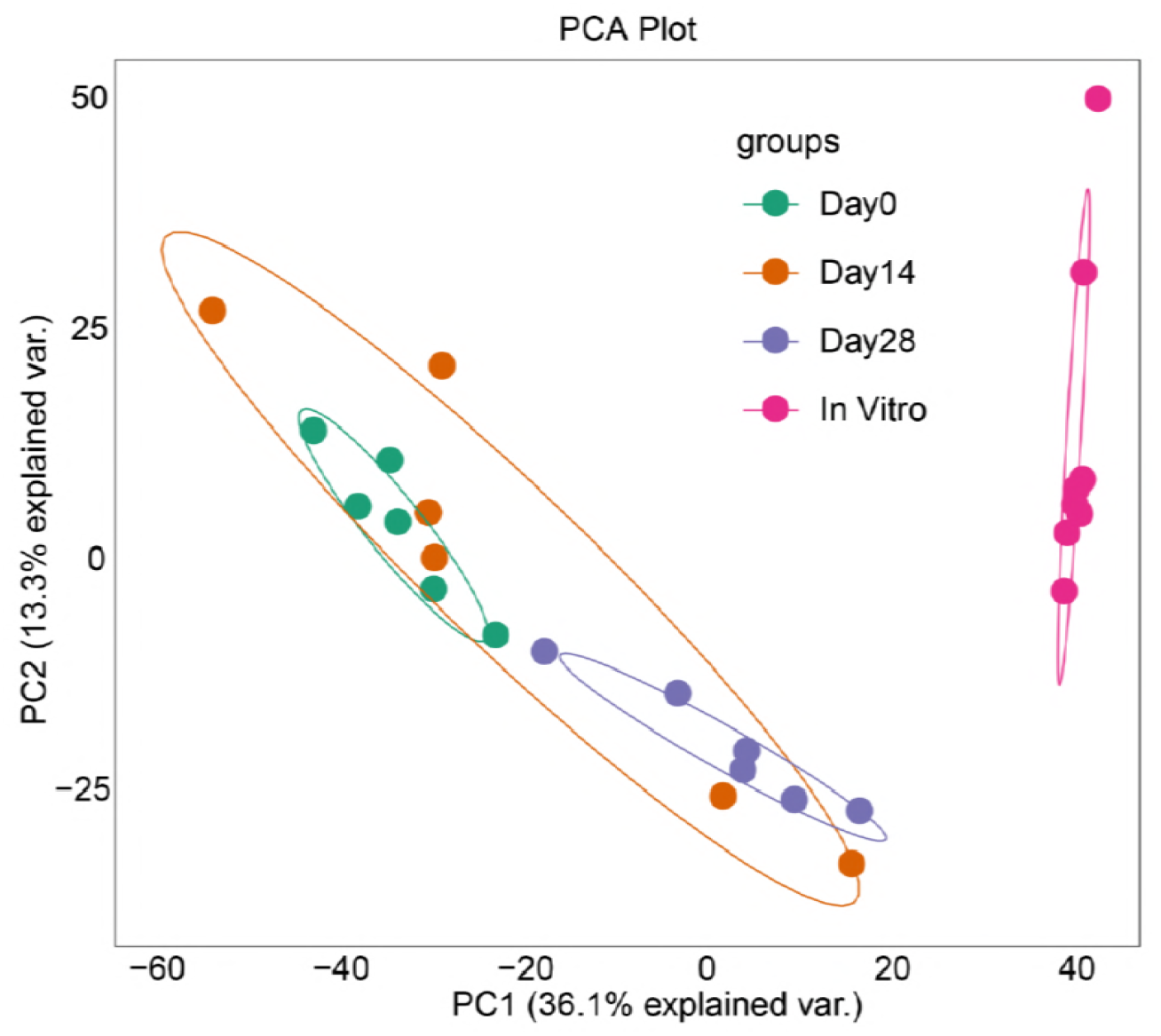
Gene expression patterns of *L. casei/paracasei in vivo* are distinct from those *in vitro*. PCA analysis showing the distinct expression pattern of *in vitro* samples compared with *in vivo* samples.

Plot of the expression correlation between any two samples showed two major convergent clusters and three minor divergent cluster (Figure S1). The largest major cluster was composed of all *in vitro* grown samples, and the second major cluster was composed of most day-28 fecal samples. One minor cluster was composed of 4 day-0 fecal samples. The other day-0 and all day-14 samples were highly convergent, constituting the other two minor clusters and suggesting a highly divergent fates of the ingested *L. casei* Zhang in different individuals, at day-14 of ingestion. These results were consistent with those of the PCA plot (Figure S1).

### The transcription modules and dynamics of gut *L. casei/paracasi* genes differentially expressed upon probiotic ingestion

To explore the transcription dynamics of *L. casei/paracasei* stain in human gut in response to the *L. casei* ingestion, we applied edgeR to compute the differentially expressed genes (DEGs) among three groups of metatranscriptomes at day-0, day-14 and day-28. A total of 1091 such DEGs were obtained using a cut off of fold change >=2 and *p*-value =<0.01. To reveal the transcription patterns of *L. casei/paracasi* DEGs in human gut, WGCNA (weighted gene co-expression network analysis) was used to analyze their expression correlation network. Two major (turquoise and blue) and three minor (green, yellow and brown) expression modules were resulted (Figure 4A).

To visualize the transcriptional dynamics of these DEG modules, we plotted their dynamic expression patterns using RPKM (reads per kilobase per million total reads) of each gene as input. Heatmap plot showed that the expression pattern of all these DEGs was similar in all six individuals prior to the ingestion of *L. casei* Zhang (day-0) (Figure 4B). The pattern at day-28 after the ingestion was also similar to each other but dramatically different from that of day-0. However, the transcription patterns were quite divergent at day-14 after the ingestion, among which four were more similar to the pattern of day-0 and two to that of day-28 (Figure 4). The heatmap dynamics well captured the PCA analysis results are shown in Figure 3. Meanwhile, this heatmap dynamics showed that the brown, yellow and green modules largely represented the individual-specific expression clusters.

### The transcription modules of *the vivo* DEGs and their associated functional clusters

We further explored the transcription patterns of their major co-expression modules using eigengene values. The module eigengene E value can be considered as a representative of the gene expression profiles in a module. Eigengene pattern of turquoise module (450 genes) showed that genes in this module were better expressed among all six day-0 samples and four day-14 samples, when compared to the very low level in day-28 samples (Figure 5A). The expression value in day-0 of individual B was higher than other day-0 individuals. The expression in three of four day-14 samples were generally higher than their corresponding day-0 samples of the same individuals (Figure 5A, upper panel). This expression pattern was varied similar to those reflected by heatmap profiling the RPKM expression values of all DEGs (Figure 4B). These observations suggested that genes in turquoise module represented the resident *L. casei/paracasei* expression, which might be transiently stimulated by the ingested *L. casei Z*hang at early time (day-14) but repressed at later time (day-28).

**Figure 4.**
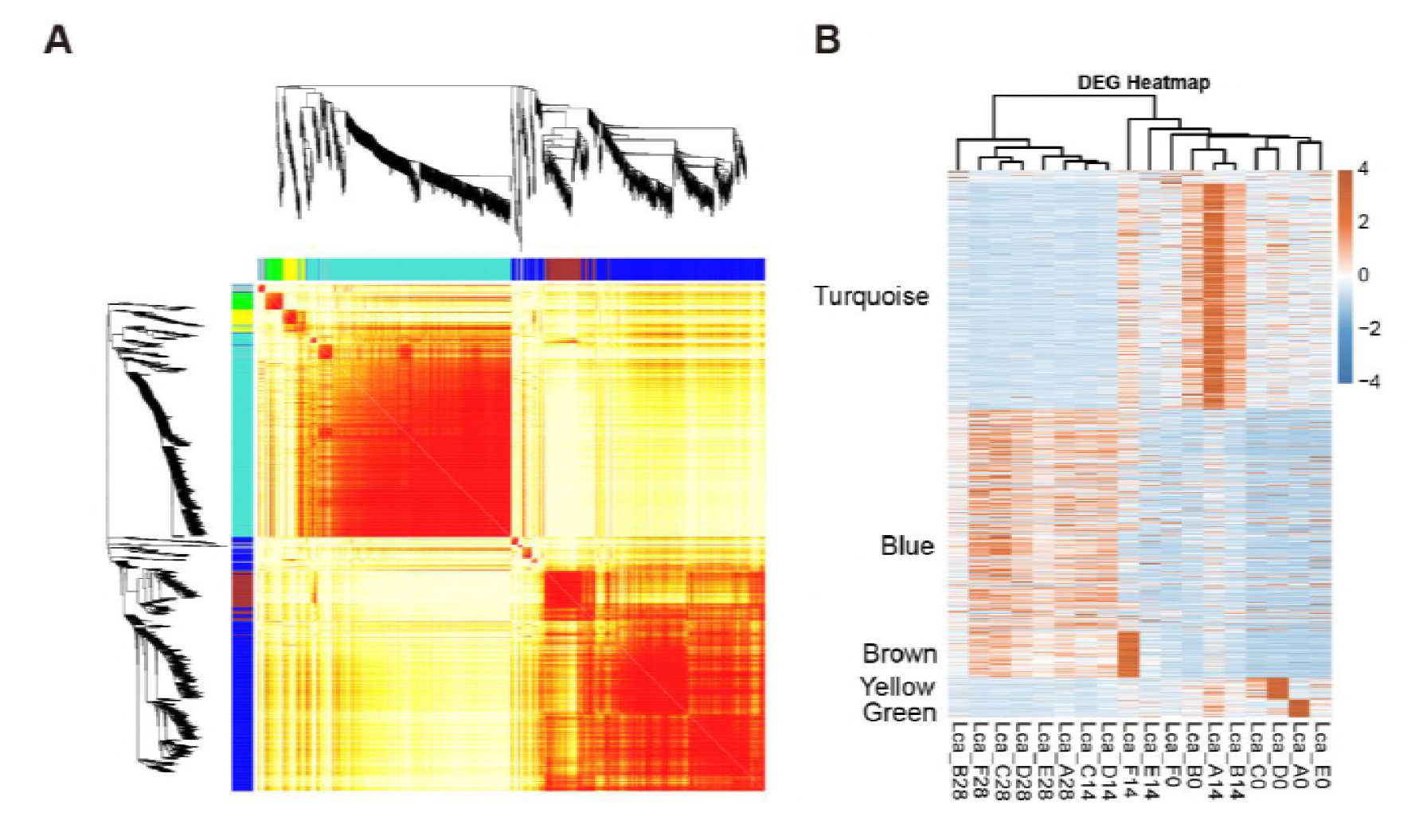
The transcriptional dynamics of differentially expressed genes in human gut upon probiotic ingestion. (A) Clustering heatmap showing the dynamical expression patterns of genes in different modules. (B) Heatmap showing the expression pattern of genes in *in vivo* samples. The genes were sorted by clustering modules.

**Figure 5.**
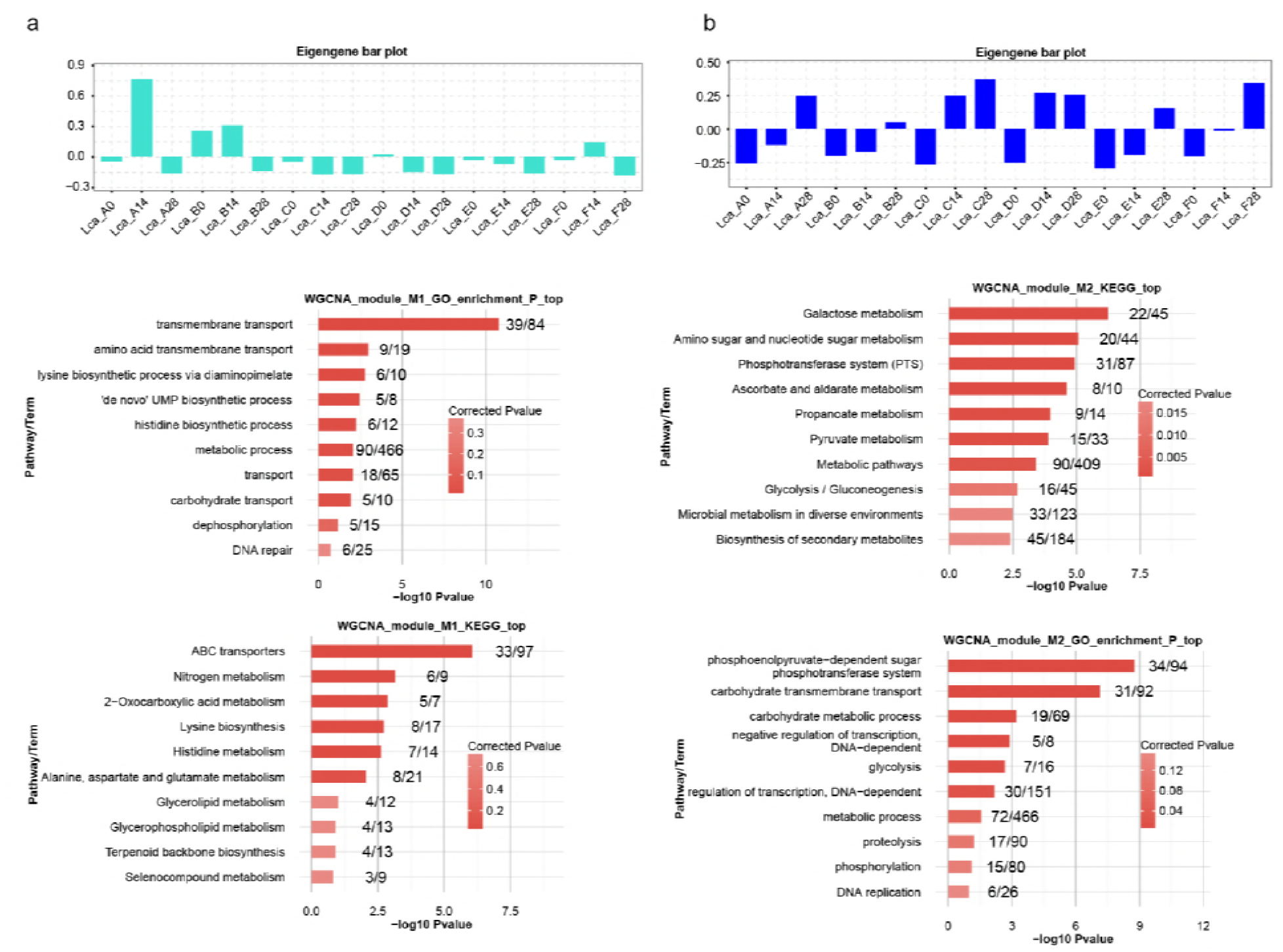
Expression pattern and functional analysis of two modules obtained by WGCNA analysis. (A) The top panel is the bar plot of eigengene value for turquoise module (first module); the middle panel is the bar plot showing the enriched GO biological processes of turquoise module; the bottom panel is the bar plot showing the enriched KEGG pathways of turquoise module. (B) The same with (A) but for the blue module (second module).

GO functional analysis showed that genes in turquoise module were enriched in transmembrane transport (*p*-value, 7.92e-12) (Figure 5A, middle panel). Genes in amino acid transmembrane transport and carbohydrate transport were enriched as well. KEGG analysis indicated that most transmembrane transport genes were ABC transporters (Figure 5A bottom panel). Another class of function more expressed by the resident *L. casei/paracasei* strains were metabolic genes (Figure 5A bottom panel), indicating that the metabolic function of the resident *L. casei/paracasei* could be altered upon probiotic expression.

Eigengene expression pattern of blue module (444 genes) showed specific expression among all six day-28 samples and 2 day-14 samples (individuals C and D) (Figure 5B), indicating that these genes were either induced by or specifically expressed from the ingested *L. casei Z*hang. Functional clustering analysis showed that these genes were enriched in sugar metabolism and transport functions including phosphoenolpyruvate-dependent sugar phosphotransferase system (GO, 34 genes, *p*-value =<1.78e-9), carbohydrate transmembrane transport (GO, 31 genes, *p*-value =<7.22e-8) and Galactose metabolism (KEGG, 22 genes, *p*-value =<2.64e-5) (Figure C middle and bottom).

Eigengene expression pattern of brown module (92 genes) were similar to that of blue module, with a major difference in gene expression pattern for individual F at day-14. In addition to the sugar metabolic function, Brown module genes were mostly enriched in ribosome and translational function (Figure S2)

### Comparison of the genome expression patterns of *L. casei in vitro* and in human gut

We next compared the transcriptome of *L. casei in vitro* and in human gut. We were aware the *in vivo* expression of *L. casei* was mixed by a fraction of resident *L. casei/paracasei.* Differentially expressed genes were obtained inside of the *in vivo* or *in vitro* groups, as well as between the *in vivo* and *in vitro* groups, which were subjected to WGCNA network analysis. Almost all *L. casei* genes (97.22%; 2871/2953) were subjected to the transcriptional regulation during *in vitro* and *in vivo* growth of the probiotic (Figure 6a). It demonstrated that M1 module contained 948 genes, representing 32.1% of all *L. casei* genes, expressed very well when grown *in vitro*, but strongly repressed when grown in human gut. These genes were expressed at relative higher level in two day-14 samples (individuals A and F), which could reflect transcripts from the transiently passed *L. casei* after being ingested. M1 module genes were enriched in KEGG pathways for translation and replication (Figure 6b).

**Figure 6.**
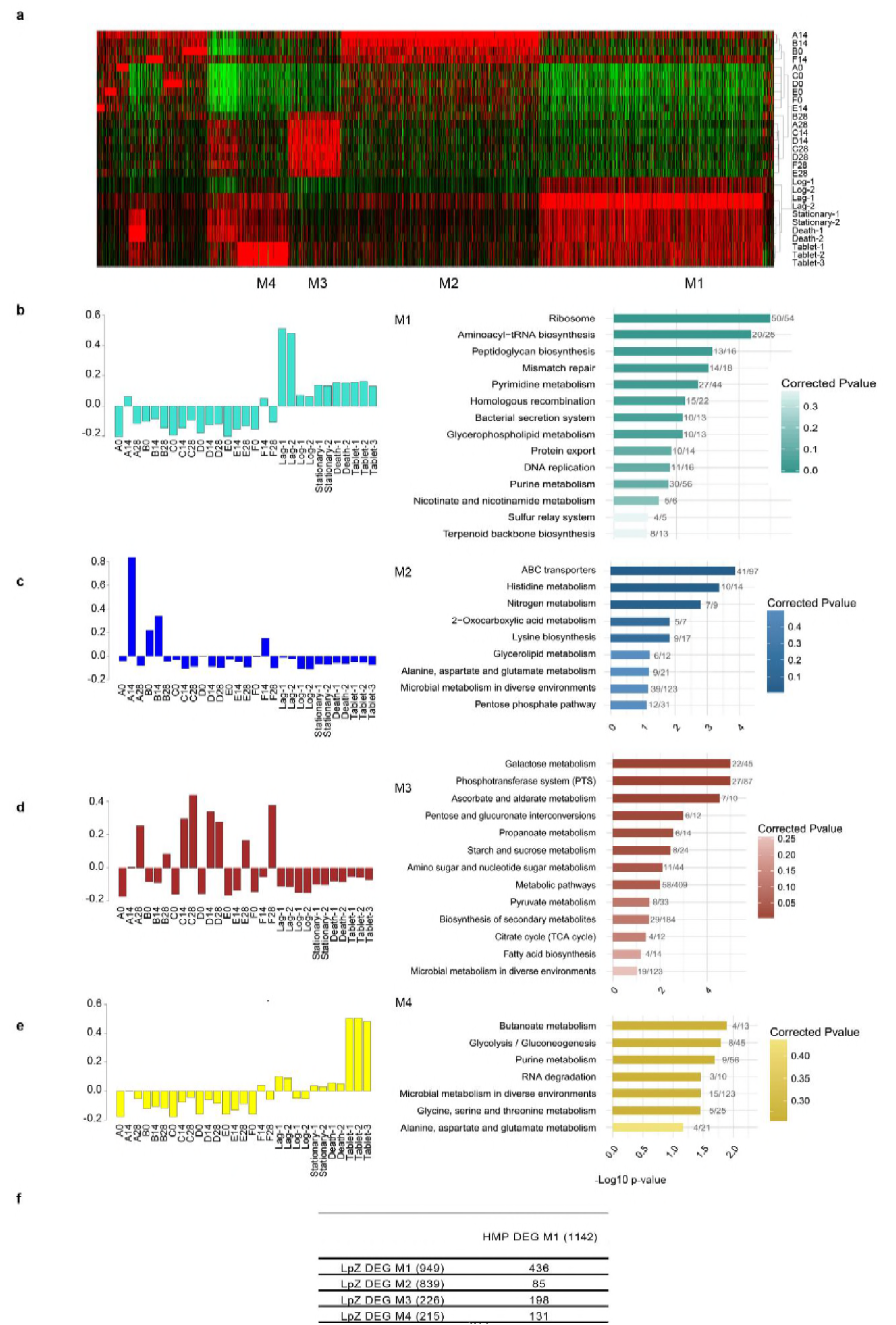
Transcriptional dynamics of the ingested *L. casei* Zhang in human gut. **(A)** Heatmap representation of differentially expressed genes mapped onto the *L. casei* Zhang genome, ranked by the co-expression modules. **(B-E)** Bar plots of eigengene value and KEGG pathway enrichment of corresponding genes in module 1 (**b**), module 2 (**c**), module 3 (**d**), and module 4 (**e**). **(F)** The overlapping genes between WGCNA results from *L. casei* Zhang and HMP mapping result.

Expression of *L. casei* Zhang genes in M2 modules (839) was induced at day-14 of three samples. These genes were strongly enriched in ABC transporters and metabolism pathways of multiple amino acids (Figure 6c), suggesting the possible presence of a transition stage, during which the ingested *L. casei* Zhang has to alter its uptake function to adapt the human gut environment. Genes in M2 modules were highly overlapped with the genes in turquoise module shown in Figure 5a.

At day-28, the late stage of ingestion, expression of a cluster of genes (226) was specifically increased (M3 module). These genes were involved in the biosynthesis and/or metabolism of the well-known probiotic molecular including galactose (20), carbohydrate utilization (33) and metabolism of propanoate the key member of SCFA (34) (Figure 6d). We found *L. casei* genes for ascorbate and aldarate metabolism were globally upregulated, suggesting a novel class of probiotic molecule. Genes in M4 module (215) were mostly expressed in the tablet form of *L. casei*, and their level in human gut was increased at the late stage of ingestion (Figure 5e). M4 genes were most strongly enriched in the metabolism of butanoate, another key member of SCFA synthesis (35).

### Dynamic expression of sRNA genes of *L. casei* Zhang *in vitro* and *in vivo*

Given the regulatory function of bacterial sRNAs(36), we then studied the possible contribution of sRNA to the highly dynamic transcriptome of *L. casei* Zhang. A total of 208 candidate sRNAs were identified from the *in vitro* grown cells. Among these candidate sRNAs, 76 were identified from all 4 stages and 143 were identified from at least two growth stages (Figure 7a). Heatmap plot of the expression patterns of all sRNAs under the *in vitro* growing states showed that although most sRNAs were expressed at multiple growth conditions, stage-specific expression of sRNAs were prevalent for *L. casei* (Figure 7b). Lag-phase, log-phase, death-phase, and tablet-phase sRNA clusters were highly specific (Figure 7b). Interestingly, stationary phase did not contain its-specific sRNA, and it rather expressed sRNA specific for the log and death phases at relatively high levels (Figure 7b).

**Figure 7.**
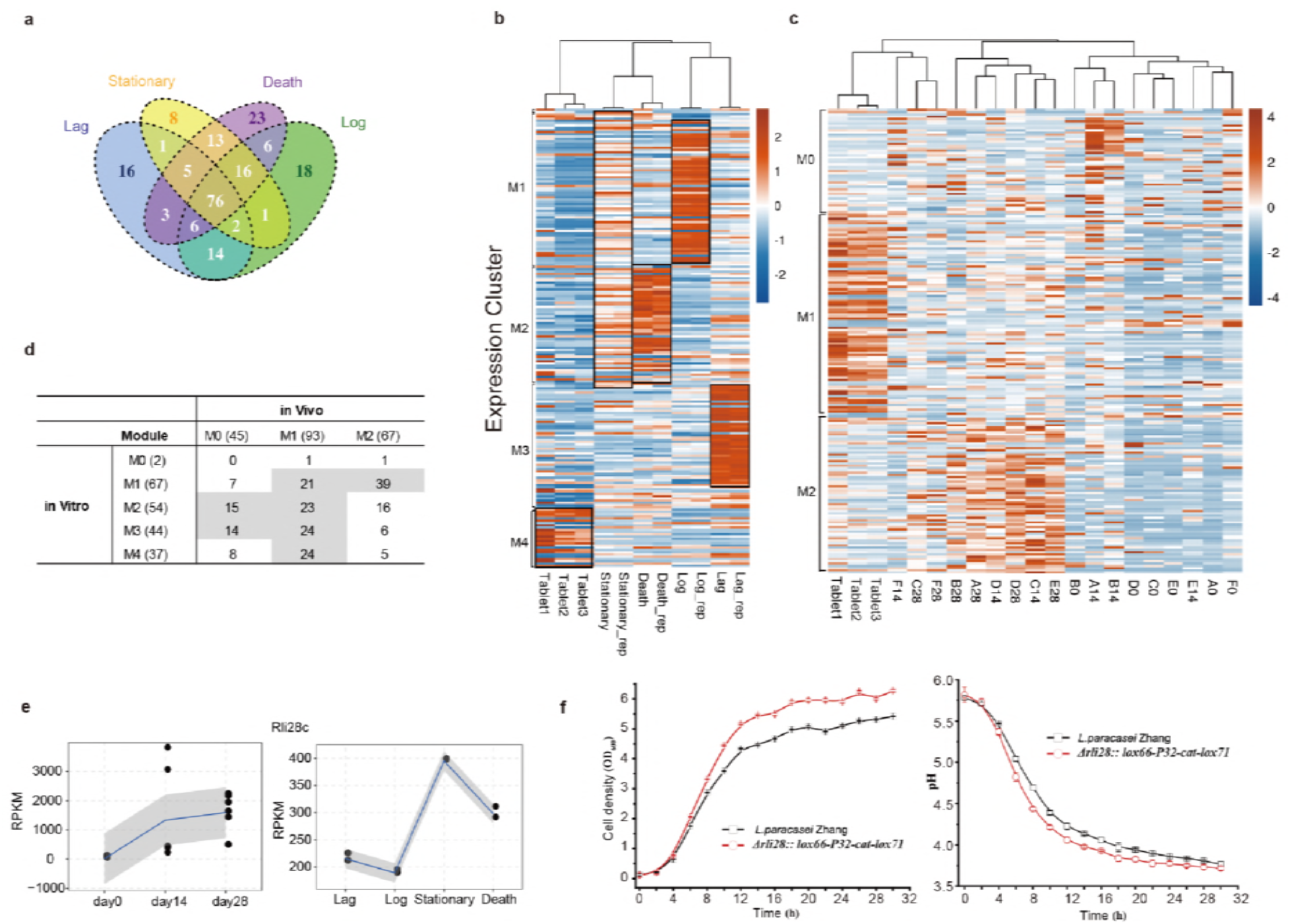
Expression profile of sRNAs and the function of rli28c sRNA. **(A)** Venn diagram showed the sRNA detection overlap among the four growth stages *in vitro*. **(B)** Heatmap presentation of the expression pattern for the *in vitro* and tablets samples by WGCNA clustering. Black rectangle represents the highly expressed sRNAs in the corresponding samples. **(C)** The same with (B) but for the *in vivo* and tablets samples. **(D)** The overlapped sRNAs numbers for major modules classified by WGCNA for *in vitro* and *in vivo* shown in (b) and (c). **(E)** The expression level line plot of RPKM values for rli28c sRNAs in *in vivo* and *in vitro* samples, respectively. **(F)** The cell density (left) and pH value of the growth medium (right) plot by time with (red) and without (black) the rli28c sRNA knockout.

When *L. casei/paracasei* was expressed in human gut, expression of sRNAs was clearly separated into two clusters. The M1 sRNAs decreased their expression after the ingestion while the M2 sRNAs increased their expression, in comparison with the sRNA expression in the tablets (Figure 7c). The *in vivo* M1 sRNAs contained sRNAs specifically expressed at each of the four in vitro grown stages at an unbiased frequency, while M2 sRNAs were mainly those of the *in vitro* log phase sRNAs (Figure 7c). This observation suggested that the *in vivo* growing state of *L. casei/paracasei* might resemble the *in vitro* log phase.

Rli28 is a small RNA that is detected in *Listeria monocytogenes* grown in stationary phase and in the intestinal lumen of its infected mice and proposed to be involved in the bacterial virulence (37). We identified five copies of rli28 expressed from the genome of *L. casei* Zhang, ranging from 210 bp to 492 bp and located in two separated loci (Table S2). We plotted the levels of rli28 genes in the *in vitro*-grown *L. casei* Zhang and the *L. casei/paracasei* grown in human gut varied greatly (Fig. 7e; Figure S3). The *in vitro* expression patterns of these rli28 genes of *L. casei* Zhang differed significantly, with one being peaked at the log phase (rli28e), two at the stationary phase (rli28c and rli28d), and two at both the stationary and death phases (rli28a and rli28b). Expression of four rli28 genes in human gut was constantly increased with the ingested time, while rli28a gene was decreased in its expression at day-28.

Rli28c peaked at the stationary phase was chosen for further functional analysis (Fig. 7f). After rli28c being knocked out using Cre/LoxP cassette, the *in vitro* growth of the mutant *LcZ* was enhanced compared to the wild-type (Fig. 7f). Meanwhile, the growth medium pH of the mutant *LcZ* was lower than the wild-type, consistent with an enhanced release of the lactic acid. Taken together, these results suggested that the stationary phase rli28c may repress the growth and the production of lactic acid by *L. casei*.

## Discussion

Exploring the fate of ingested probiotics thoroughly at transcriptional level remains challenge thus far. To our best knowledge, this study presented the first effort to profile the transcription of mRNA and sRNA of probiotic and resident *Lactobacillus* in human gut, by extracting the transcriptome reads of the probiotic bacteria from metatranscriptomic reads. Classical metatranscriptomic analysis shows that the ingested probiotic bacteria does not alter the global composition of gut microbial community to any appreciable level compared to the individual variations, consistent with the previous results (31, 32, 38-40). Comprehensive comparative transcriptome analysis was performed in human fecal samples at different time points of ingestion, and between the *in vivo* and *in vitro* growth states. These comparative transcriptomic and metatranscriptomic studies led to some interesting findings.

### The resident *L. casei/paracasei* strains transcribe differently from the ingested probiotic *L. casei* strain

It has been nearly impossible to study the transcriptome of individual strains among the large microbial community in human gut previously (41). In this study, we have used deep sequencing technology to obtain metatranscriptomes of human gut microbiota from the fecal samples of six healthy volunteers, followed by mapping the metatranscriptomic reads onto the genomes of the ingested *L. casei* Zhang and its two close relatives *L. casei/paracasei* strains. This approach allowed us to compare the transcription of the ingested *L. casei in vitro* and its close relatives in human gut (day-0 samples), demonstrating that transcription of both mRNAs and sRNAs differ greatly *in vitro* and *in vivo*.

We have also applied strain-specific reads to distinguish the transcripts from three closely related *L. casei/paracasei* strains, indicating that the increased mapping of metatranscriptomic reads is primarily derived from the ingested *L. casei*. We proposed that a combination of increased metatranscriptomic sequencing depth and stain-specific mapping strategy might allow a higher resolution of the transcriptomes of various microbe strains in human and other mammals, as well as the transcriptome dynamics in response to the exogenous bacteria, in the future.

### The fate of the ingested *L. casei*: death/lysis or changing the transcription pattern

Probiotic microorganisms can generally survive well when they pass through the stressful GI tract conditions in a few hours, and stay in colon for a few days (14, 15, 25). Microbial cells that cannot survive the GI tract undergo cell lysis (14, 42). It is unclear what is going on at the transcriptome level when the probiotics were ingested. In this study, we have demonstrated that transcription of the ingested *L. casei* does not inherit the *in vitro* transcription pattern at all. Moreover, transcription patterns at day-14 and day-28 differ significantly.

These findings have an important implication regarding the fate of ingested bacteria and what we are detecting from the fecal samples. It is generally worried that during the course of probiotic uptake, the majority of probiotics that we detected from fecal samples are the dead bacteria after being ingested. However, the distinct transcription patterns between *in vitro* and *in vivo*, as well as between those after 14 days and 28 days of probiotic uptake, strongly suggest that the detected probiotic transcriptomes reflect those have survived the GI tracts. Our results support the previous hypothesis of the cell lysis for the dead ingested bacteria (20, 42). We do not exclude the possibility that the dead probiotic bacteria might still yield fragmented DNA signals. However, the dead probiotic *L casei* unlikely yield RNA signals according to our reported transcription patterns. In conclusion, this study suggests that transcriptome analysis represents a more effect way for detecting the living bacteria in fecal samples.

### Activation of ABC reporters might be required for probiotic survival during the early stage of ingestion

ATP-binding cassette (ABC) transporters represent one of the largest classes of transporters using the power from ATP hydrolysis to drive the translocation of different substrates across cell membranes (43). ABC transporters not only transport a large variety of nutrients into cells from environments, but also transport various cellular components away from the cells. For example, multidrug ABC transporters transport a wide range of drugs from cell (44). In this study, we found that in three day-14 and one day-0 fecal samples, genes encoding ABC reporters were globally activated in *L. casei*, compared with their expression under *in vitro* growth condition. As we have shown, upon *L. casei* Zhang ingestion, the increased *L. casei* mapping is from the ingested *L. casei*. The increased expression of ABC transporters should therefore indicate that the *L. casei* Zhang survived GI tracts has changed its expression pattern favoring the expression of ABC transporters. Activation of the expression of ABC transporters might enhance the ability of ingested *L casei* in uptaking of nutrient from the human gut environment.

It is known that human gut microbes establish direct chemical interactions with host (45). It could be possible that the signals for the global activation of ABC transporters were sent by the gut microbial community, reflecting its early cross-talk with the ingested probiotic. On the other hand, activation of ABC transporters could also reflect how the ingested *L casei* respond to the living condition in human gut.

### Genes for sugar and SCFA metabolisms are activated during the later stage of probiotic ingestion

Interestingly, we observed a clear shift of transcriptional patterns between day-14 and day-28 samples, in which the activated expression of ABC transporter disappeared and activated expression of genes for galactose and sugar metabolism appeared. This shift indicates a dynamic cross-talk between ingested *L. casei* and human gut microbiota. It could be possible that the early cross-talk elicits a signal for activated expression of ABC reporters. However, as *L. casei* uptake continues, the interaction between the ingested *L. casei* and human gut microbes has been established, the signal for activation of ABC transporters of the ingested *L. casei* might then lose. Instead, signals for galactose and sugar metabolism are secreted, be sensed and reacted by the ingested *L. casei*.

Human gut microbiome is developed with its host after birth, which modulates the host metabolic phenotype(45). The host and microbiome establish metabolic axes resulting in combinatorial metabolism of substrates by the microbiome and host genome, which produce various metabolites such as bile acids, choline, and short-chain fatty acids (SCFAs) that are essential for host health (46, 47). It is interesting to observe that at the later stage of the ingestion of probiotic *L. casei* genes for galactose and sugar metabolism, as well as those for the metabolism of one class of SCFAs propanoate (48, 49) were globally activated. These findings are consistent with the current knowledge that probiotic bacteria can contribute metabolites such as acetate, lactate and propanoate (14, 50, 51). A number of reports have shown that Lactobacillus stains produce SCFAs (52, 53). The increase in propionic acid is dependent on the intake time, much more pronounced after 3 weeks of intake than after eight days, which agrees well with our observed time-dependent activation of genes for propanoate metabolism.

### The highly regulated expression of *L casei* sRNAs and growth repression by sRNA *rli28*

Small RNAs represent a large class of novel regulatory molecules in bacteria (36, 54). The sRNAs in *Lactobacillus* have not been well characterized before. In this study, we have identified 208 sRNAs in *L. casei* Zhang growing under four different growth stages *in vitro*, among which 76 were all overlapped. Almost all sRNAs display a stage-specific growth pattern, which agrees well with the regulatory roles of sRNAs (55, 56).

After the intake, we found that sRNAs highly expressed in the death and stationary phases were well expressed in human gut. By creating a lox knock-out *L. casei* Zhang, we have shown that one copy of rli28 who best expressed in stationary phase inhibits *L. casei* growth *in vitro*. This suggests that sRNAs could regulate the bacterial growth rate.

## Conclusions

Study the transcription of the ingested probiotic in human gut using the metatranscriptome profiling approach has shown that the probiotic strain transcribes in a unique way different from its *in vitro* transcription and the *in vivo* transcription of the closely related species. Expression of about 40% of mRNAs and sRNAs is repressed, while genes encoding ABC transporters and those in sugar and SCFA metabolisms are activated at the early and later stages of ingestion, respectively. The unique transcription pattern of the probiotic bacteria *in vivo* might shape their characteristics of being transient passenger without much affecting of the resident gut microbiota. These findings together underline the presence of a dynamic crosstalk between the probiotic and human gut including the microbial community, which ensures a tightly regulated expression of the probiotic genome in *vivo*, which are worth of further studies in the future. Moreover, the developed methodology can be extended to study the *in vivo* expression of probiotics and pathogens.

## Methods

### Subjects and study design

Subjects were asked to orally intake 4 probiotic tablets consisting of a total of 10.6 Log_10_ CFU *L. casei* Zhang daily from Day 0 to 28. Fecal samples were collected from the subjects on Days 0, 14 and 28 in sterile containers and were kept refrigerated. Samples were transported on ice to the laboratory within 2 hours, and were kept at -80°C until further analysis.

### Stool collection, storage, fecal RNA extraction and sequencing

Stool samples were respectively collected before and after a 4-week consumption period. Gut microbiota were sampled by non-invasively fecal collection. Stool samples were taken in duplicate by coring out feces with inverted sterile 1 mL pipette tips. These tips were then deposited in 15 mL Falcon tubes. Samples collected at home were stored temporarily at −20°C until transported to the laboratory and then stored in −80°C freezers. Subject samples collected abroad were stored at −20°C, shipped to the company on dry ice, and then stored at −80°C. Total RNAs were treated with RQ1 DNase (promega) to remove DNA. The quality and quantity of the purified RNA were determined by measuring the absorbance at 260 nm/280 nm (A260/A280) using smartspec plus (BioRad). RNA integrity was further verified by 1.5% Agrose gel electrophoresis. For each sample, 5 μg of total RNA was used for RNA-seq library preparation. Ribosomal RNAs were depleted with Ribo-Zero™ rRNA depletion kit (Epicentre, MRZB12424) before used for directional RNA-seq library preparation (gnomegen K02421-T). Purified mRNAs were iron fragmented at 95°C followed by end repair and 5’ adaptor ligation. Then reverse transcription was performed with RT primer harboring 3’ adaptor sequence and randomized hexamer. The cDNAs were purified and PCR amplified. PCR products corresponding to 200-500 bps were purified, quantified and stored at -80 °C until used for sequencing.

### *In vitro* sample RNA extraction, library construction and sequencing

For the *in-vitro* bacterial samples, we collected the samples by two different styles. As for the first style, we cultured the *L. casei* Zhang on the medium and collected two replicate samples from each of the four growth stages, lag, log, stationary, and death stage, respectively. For the second, we collected the samples from the probiotic tablets same as the above, and three replicates were prepared. After sample collection, total RNAs were extracted from samples mentioned above by using Trizol Reagent (Invitrogen). Then we used Ribo-Zero rRNA removal kit to remove the rRNAs. After that, extracted RNA was amplified using custom barcoded primers and sequenced with paired-end 100 bp reads by Illumina HiSeq2500 platform.

### Quality filtering and sequence statistics

After sequencing, raw reads would be first discarded if containing more than 2-N bases, then reads were processed by clipping adaptor, removing low quality reads and bases from the end of each reads and discarding too short reads (less than 16nt) by FASTX-Toolkit (Version 0.0.13). The metagenomic, metatranscriptomic and the *in vitro* samples were filtered with the same method and parameters.

### Data validation by qPCR

Genomic DNA and total RNA were extracted from fecal samples of each volunteer. To validate genes copy number from metagenomic sequencing, quantitative Polymerase Chain Reaction (qPCR) was applied to detect the relative copy numbers using ABI Prism 7300 Real-Time PCR System with standard procedures. A known fragment, containing 3’-UTR of RORA gene (human) was inserted into psiCHECK 2 plasmid. The plasmid was added into each sample by quantitation, and detected as an external control by specific primers. The relative level of DNA level was analyzed after being normalized by the external control.

For metatranscriptomic mRNA detection, total RNAs was extracted from the same fecal samples of each volunteer for sequencing. To ensure there was no genome DNA contamination, RNA was treated with DNAse 1 (Takara) for 2h, and then applied to PCR validation. The mRNA fragments of β-actin (human) obtained by in vitro Transcription (Transcript Aid T7 High Yield Transcription Kit, Thermo Scientific) was added into each RNA samples and applied to the reverse-transcribed by random hexamer primers using M-MLV reverse transcriptase (Promega). RT-qPCR was performed using ABI Prism 7300 Real-Time PCR System with standard procedure, and the relative expression level of genes were normalized by β-actin. The PCR primers were provided in Table S3.

### HMP database retrieval

We chose HMP database (http://hmpdacc.org/) as reference to do the structural and functional analysis. First, we downloaded the complete genome sequences and annotation of human gut microbiome, which contains 358 publicly available human microbiome genomes generated from the National Institutes of Health (NIH) Human Microbiome Project and the European MetaHIT consortium. Besides, we added the *L. casei Zhang* genome (http://www.ncbi.nlm.nih.gov/) to the database to evaluate the influence of *L. casei Zhang* to the microbiome. We then aligned our metagenomic and metatranscriptomic data to the genomes with bowtie2(57), allowing no more than one mismatch. To deal with cases of multiple mapping, we selected no more than 10 best matches of the alignment based on the mapping quality, and then we divided the reads by its hits number, and each hit occupied one part of the reads. After that, we calculated the reads number and RPKM value for each contig and gene in the database. We then obtained the abundance of different taxonomic levels from species to kingdom by adding relative contigs abundance together. To consistently estimate the functional composition of the samples, we annotated the genes from the HMP database using COG orthologous groups and KEGG pathways by blastx program with e-value 1e-5. We ensured that comparative analysis using these procedures was not biased by data-set origin, sample preparation, sequencing technology and quality filtering.

For metatranscriptomic gene abundance, to study gene expression alteration changed by the *L. casei Zhang*, we compared the expression change between day 14 and day 0, day 28 and day 0 and day 28 and day 14. First, we got differentially expressed species and extracted all genes abundance from these species, and then obtained the differentially expressed genes. We then used WGCNA(58) method to classify the differentially expressed genes as modules based on their expression pattern. After classification, we used the annotation of KEGG to obtain the functional enrichment pathways by hypergeometric test.

### *In vivo* and *in vitro* samples co-analysis

To find the transcriptome difference of *L. casei Zhang* between *in-vivo* and *in-vitro* samples, we compared the gene expression difference among these samples by aligning the transcriptome reads to the *L. casei Zhang* genome. We used bowtie2(57) software to align reads to the *L. casei Zhang* genome allowing 1 seed mismatch. RPKM value for each gene was calculated for each sample. Then we compared the gene expression changes between each samples groups with each other by edgeR (59) package. Samples *in-vivo* of each point was compared with samples *in-vitro* of each stage and type, and samples *in-vivo* was compared with each other, samples *in-vitro* was compared with each other. We then used WGCNA(58) method to classify the differentially expressed genes as modules based on their expression pattern. After classification, we used the annotation of KEGG to obtain the functional enrichment pathways by hypergeometric test.

### Bacteria sRNA prediction and expression analysis

To have an exact prediction of *L. casei* Zhang sRNAs, we developed an algorithm to detect peaks from alignment results among intragenic, intergenic (between two adjacent genes) and antisense regions. We used the RNA-seq data from four stage bacterial strain cultured on the medium. We merged the mapping result file from the same stage, and ran the computer program separately for the four stages. After prediction, we merged the sRNAs predicted from the four stages by genomic locations and got a final sRNA prediction result. The detail description of algorithm is described below. Based on the alignment result, 5 bp window size was chosen as the default window size. Peak starting site was identified as the end of one window, the median depth of which is no more than 0.25 fold of all of the adjacent downstream eight windows. Peak terminal site was identified as the start of one window whose median depth is no more than 0.25 fold of all of the adjacent upstream eight windows. After the algorithm realization, we then filtered the peaks according to the following three thresholds: 1) the length of peaks should range from 40bp to 500bp; 2) the maximum height of one peak should be no less than 60 read depth; 3) the medium height of one peak should be no less than 20 depth. After peak definition, we classified the peaks into three different classes according to their locations: 1) intragenic peaks whose locus were overlapped with known mRNA genes and on the same strand; 2) antisense peaks were defined as peaks whose locus were overlapped with known mRNA genes but on the opposite strand; 3) intergenic peaks whose locus were neither overlapped with known mRNAs on the same strand nor on the opposite strand. Antisense and intergenic peaks were defined as sRNAs. We aligned the sRNA sequence to the Rfam database (version 12.0) (60) to identify homologies from related bacteria by Blast method (E-value ≤ 1e-5).

After sRNA prediction, we got the normalized expression level of each sRNA for each samples. We then used WGCNA(58) method to classify the differentially expressed sRNAs as modules based on their expression pattern.

### sRNA knockout experiment

To validate the influence on bacteria by sRNAs, we selected sRNAs that expressed significantly and dynamically to do the knockout experiment. Rli28 and ratA from the plasmid of *L. casei Zhang* were chosen. The target sequence of Rli28 is TTAATGCGATTAAAGCCACGGTAAAGGTACCGAAAGCCAGCATTAATTGTAAAGCGTCCGCAACGGACACTTAGGCTACTCCTTTCATTAGGATTTATGGGCTTTAGGGGTTTAACACCATAAGCACCACCTCCGATCGGAAATAGCCACCGCCTTAACTTCTCTACAAGCTTTAATTATACAGGAGCTTT, which locates on the plasmid from 30466 to 30656. The target sequence of ratA is TAATATAGACAGAAAAAGGGAAGCCCCGCTAGAACAGGACTTCCCATGCAAGCCGCTTCAAAGGCGGTGGCAGAAATTTAATAAACGATTTT, which locates on the plasmid from 28019 to 28110. The knockout experiment was performed according to one published protocol for gene deletions in *Lactobacillus*(61), and the knockout efficiency of Rli28 was validated by RT-PCR. After knockout, we tested the cell density and pH levels of the knockout bacteria with three independent replicates.

### MetaPhlAn2 analysis

For both metagenomic and metatranscriptomic reads, we have applied the MetaPhlAn2 and GraPhlAn software(62) to obtain the relative abundance of each species. Top abundant species of all samples were used to make a dendrogram heatmap via hierarchical clustering. After the calculation of species abundance, we got differentially expressed species to analysis the influence of *L. casei Zhang* on transcription variation.

### Other statistical methods

Principle Component Analysis (PCA) was used to analyze the time and individual influence. Fisher Exact Test was used to obtain the enrichment of each functional cluster. Statistical figures and tables were obtained by a free statistical software R. Cluster was performed by the Cluster3.0 software and the heatmap was generated by Java TreeView (http://bonsai.hgc.jp/∼mdehoon/software/cluster/software.htm).

## Abbreviations

GALT: gut-associated lymphoid tissue
IBD: inflammatory bowel disease
IBS: irritable bowel syndrome
*L. casei* Zhang: *Lactobacillus paracasei* Zhang
ORFs: Open Reading Frames
PCA: Principal Component Analysis
PTR: peak-to-trough ratio
qPCR: quantitative Polymerase Chain Reaction
RPKM: Reads Per million per kilobase
SCFAs: short chain fatty acids

## Declarations

### Ethics approval and consent to participate

The experiment was approved by the Ethics Committee of the Inner Mongolia Agricultural University (Hohhot, China). A written consent was obtained from every volunteer.

### Consent for publication

All volunteers participated in this paper have signed to give the consent for publication.

### Availability of data and material

The sequences reported in this paper have been deposited in the National Center for Biotechnology Information Sequence Read Archive under accession no. SRP065752.

## Authors’ contributions

H.Z. and Y.Z. led the project; H.Z., Y.Z., J.W. and Z.S. conceived and designed the project; Y.Z., H.Z., J.Q. and D.C. wrote the manuscript; X.L., J.D. and J.Z. collected samples and performed experiments; D.C., C.C., and Q.Hou analyzed the data and generated graphics.

## Acknowledgements

We would like to express our gratitude to members in Prof. Heping Zhang’ team and the team from ABLife for their assistance in preparation of samples and sequencing libraries. We would like to thank Ms. Hong Wu (ABLife) for her help in language editing.

